# Gradual divergence and diversification of mammalian duplicate gene functions

**DOI:** 10.1101/004267

**Authors:** Raquel Assis, Doris Bachtrog

## Abstract

Gene duplication provides raw material for the evolution of functional innovation. We recently developed a phylogenetic method to classify the evolutionary processes underlying the retention and functional evolution of duplicate genes by quantifying divergence of their gene expression profiles. Here, we apply our method to pairs of duplicate genes in eight mammalian genomes, using data from 11 distinct tissues to construct spatial gene expression profiles. We find that young mammalian duplicates are often functionally conserved, and that functional divergence gradually increases with evolutionary distance between species. Examination of expression patterns in genes with conserved and new functions supports the “out-of-testes” hypothesis, in which new genes arise with testis-specific functions and acquire functions in other tissues over time. While new functions tend to be tissue-specific, there is no bias toward expression in any particular tissue. Thus, duplicate genes acquire a diversity of functions outside of the testes, possibly contributing to the origin of a multitude of complex phenotypes during mammalian evolution.

## Introduction

Gene duplication produces copies of existing genes, which can diverge from their ancestral states and contribute to the evolution of novel phenotypes. A large proportion of mammalian genes arose via gene duplication (Li et al. 2001; Ryvkin et al. 2009), and many are members of gene families with diverse and essential functions. For example, Hox, growth factor, and olfactory receptor gene families were all produced by gene duplication. However, the evolutionary paths leading from functionally redundant duplicate copies to distinct genes with important functions remain unclear.

Different processes may drive the long-term retention and functional evolution of duplicate genes: Parent and child copies may each maintain the function of their ancestral single-copy ortholog (conservation; Ohno 1970); one copy may maintain the ancestral function, while the other acquires a new function (neofunctionalization; Ohno 1970); each copy may lose part of its function, such that together both copies carry out the ancestral function (subfunctionalization; Force et al. 1999; Stoltzfus 1999); or both copies may acquire new functions (specialization; He and Zhang 2005). We recently developed a method that utilizes distances between gene expression profiles to classify these evolutionary processes (Assis and Bachtrog 2013). Our method is applied to pairs of duplicates and requires that, for each pair, we can distinguish between parent and child copies and identify a single-copy ancestral ortholog in a closely related sister species. Moreover, parent, child, and ancestral genes must all have spatial or temporal gene expression data from which gene expression profiles can be constructed.

To study the roles of conservation, neofunctionalization, subfunctionalization, and specialization in the retention of mammalian duplicate genes, we applied our method to pairs of duplicate genes from eight mammalian genomes: human (*Homo sapiens*), chimpanzee (*Pan trogodytes*), gorilla (*Gorilla gorilla*), orangutan (*Pongo pygmaeus abelii*), macaque (*Macaca mulatta*), mouse (*Mus musculus*), opossum (*Monodelphis domestica*), and platypus (*Ornithorhynchus anatinus*). Using synteny information from whole-genome alignments to determine orthologous genomic positions, and parsimony to infer the evolutionary dynamics of genes, we distinguished between parent and child copies and identified ancestral single-copy orthologs for each pair of duplicates. Then, we applied our classification method to RNA-seq data from 11 mammalian tissues: female and male cerebrum, female and male cerebellum, female and male heart, female and male kidney, female and male liver, and testis (Brawand et al. 2011).

## Results

In total, we obtained 654 pairs of mammalian duplicate genes for which we could distinguish between parent and child copies and also identify at least one expressed single-copy ancestral gene in a closely related sister species. Application of our method to these pairs yielded 382 cases of conservation, 213 cases of neofunctionalization (105 neofunctionalized parent copies, 108 neofunctionalized child copies), 9 cases of subfunctionalization, and 50 cases of specialization. Thus, most mammalian duplicate genes have conserved functions. Moreover, functional divergence typically affects only one gene copy, and retention of duplicates by subfunctionalization is rare.

Comparing duplicates from mammalian genomes of different evolutionary distances enabled us to examine whether there is a negative relationship between functional conservation and age of duplicate genes, as expected if genes evolve new functions over time. We used parsimony to date the origin of child copies along the mammalian phylogeny (Figure 1A). Consistent with global patterns, conservation is the most common evolutionary process underlying the retention of duplicate genes in every mammalian lineage examined (Figure 1A). To test if functional conservation decreases with increasing evolutionary divergence between species, we calculated rates of protein sequence divergence (*K*_a_) between single-copy genes in human and each sister species on the tree, and used these values as estimates of evolutionary divergence between pairs of species. Comparison of median *K*_a_ to proportions of duplicate gene pairs with conserved functions revealed that functional conservation of duplicates indeed decreases significantly with evolutionary divergence between species, and that this decrease is approximately linear (Figure 1B). Thus, young pairs of mammalian duplicates are generally functionally conserved, and new functions evolve gradually over time. Moreover, *K*a is not as strongly correlated to the proportion of functionally conserved single-copy genes, indicating that functional divergence occurs faster in duplicates than in single-copy genes.

**Figure 1.**
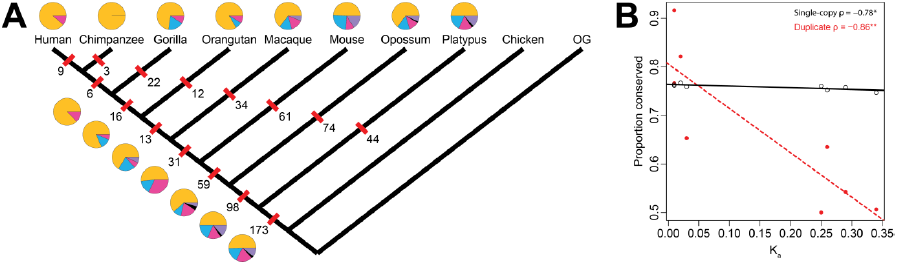
Evolutionary processes driving the retention of mammalian duplicate genes. *A*) Pie charts depicting the role of each process on different branches of the mammalian phylogeny (yellow = conserved; blue = neofunctionalization of parent copy; pink = neofunctionalization of child copy; black = subfunctionalization; purple = specialization). Numbers of duplicate gene pairs examined along each branch are indicated beside red ticks. *B*) Relationship of median *K*_a_ between pairs of species to proportions of functionally conserved single-copy genes (black) and pairs of duplicate genes (red). Linear regression lines are depicted to show rate of decreased functional conservation, and Pearson’s correlation coefficients are shown in the top right corner of the plot. * *p* < 0.05; ** *p* < 0.01.

To determine the types of novel functions acquired by mammalian duplicates over time, we examined differences between gene expression patterns in copies of pairs retained by neofunctionalization. In such cases, one copy has maintained the ancestral function (the “conserved” copy), while the other has acquired a new function (the “neofunctionalized” copy). Thus, we can directly assess ancestral and new functions within pairs. We used the highest relative tissue expression level for each gene as a measure of its tissue specificity. Comparison of distributions of tissue specificities revealed that, as expected, conserved copies and ancestral genes have similar tissue-specific expression levels (Figure 2A). Single-copy genes have similar tissue-specific levels as well, indicating that duplicate genes are initially as broadly expressed as single-copy genes in the genome. In contrast, neofunctionalized copies are significantly more tissue-specific than ancestral, conserved, and single-copy genes. An alternative metric of tissue-specific expression, τ (Yanai et al. 2005), yields the same conclusions (Figure S1). Thus, duplicates are initially as broadly expressed as typical single-copy genes, and acquisition of a new function by neofunctionalization results in increased tissue specificity.

**Figure 2.**
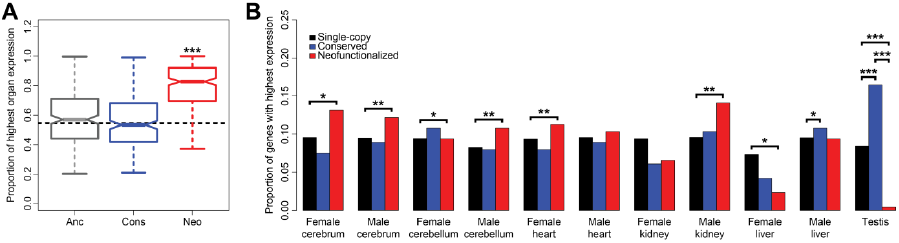
Comparison of tissue-specific expression in conserved and neofunctionalized copies of pairs that underwent neofunctionalization. *A*) Boxplots of highest relative expression levels for ancestral (Anc, gray), conserved (Cons, blue), and neofunctionalized (Neo, red) genes. Dotted black line represents median for single-copy genes, and asterisks show significance relative to distribution of single-copy genes. *B*) Barplots depicting proportions of single-copy (black), conserved (blue) and neofunctionalized (red) genes with highest expression in each tissue. Asterisks above lines connecting two bars indicate significance between classes. * *p* < 0.05; ** *p* < 0.01; *p* < 0.001.

To assess whether conserved and neofunctionalized copies of pairs show different expression patterns across tissues, we examined quantities of genes with highest expression levels in each tissue (Figure 2B). In most tissues, numbers of highly expressed single-copy genes are similar to those of conserved gene copies. The two exceptions are male liver and testis. While the difference in highly expressed male liver genes is modest, the proportion of conserved testis-specific genes is nearly double that of single-copy testis-specific genes. In contrast, only a small proportion of neofunctionalized copies are testis-specific. Thus, many genes initially arise with testis-specific functions and gradually acquire other functions over time, supporting the “out-of-testes” hypothesis of new gene origination (Kaessmann 2010). Moreover, while functions of neofunctionalized copies are typically more tissue-specific (Figure 2A), there is no bias toward specificity in any particular tissue(s). Hence, it appears that mammalian duplicate genes acquire new functions in a diversity of tissues.

## Discussion

In a recent study, we applied our classification method to pairs of duplicate genes in *Drosophila melanogaster* and *D. pseudoobscura* (Assis and Bachtrog 2013), for which the median *K*s is 1.79 (Richards et al. 2005). Contrary to our observation in mammalian duplicates, we found that most *Drosophila* duplicates were neofunctionalized, and examination of evolutionary processes over shorter divergence times suggested that novel functions arise rapidly (Assis and Bachtrog 2013). However, the smallest *K*s examined in *Drosophila* was 0.11 (between *D. melanogaster* and *D. simulans*) (Lazzaro 2005). In contrast, while the *K*s between human and platypus is approximately 1.41 (Warren et al. 2008), the smallest *K*s examined in mammals was 0.01 (between human and chimpanzee; Chen and Li 2001), which is an order of magnitude smaller than that between any pair of *Drosophila* species we analyzed (Assis and Bachtrog 2013). Thus, we have greater temporal resolution in mammals than in *Drosophila*, enabling us to more closely examine the functional diversification of mammalian duplicates over evolutionary time.

In contrast to the widespread and rapid neofunctionalization observed in *Drosophila*, young mammalian duplicates are primarily conserved, and new functions arise slowly over time. This difference may be due to the larger effective population size (*N*_e_) of *Drosophila* than of mammals (Beckenbach et al. 1993; Lynch and Conery 2003; Jensen and Bachtrog 2011), which contributes to more efficient adaptive protein sequence evolution in *Drosophila* (Britten 1986; Moriyama 1987; Carroll 2005), and could similarly result in more rapid acquisition of adaptive functions by *Drosophila* duplicate genes. Furthermore, while small *N*_e_ is also thought to result in a higher prevalence of subfunctionalization (Lynch et al. 2001), this process does not appear to play a major role in the retention of duplicate genes in either lineage. A possible reason for this observation is that subfunctionalization may be more common in duplicate genes produced by whole genome duplication events (Casneuf et al. 2006; Fares et al. 2013), which our study does not examine.

In both *Drosophila* and mammalian duplicate genes, we uncovered strong support for the “out-of-testes” hypothesis of new gene emergence (Kaessmann 2010). Testes may facilitate the initial transcription of young genes, while sheltering them from pseudogenization as they acquire new functions (Kaessmann 2010), and may thus be an ideal tissue for young genes. However, while there does not appear to be a bias in evolution of functions from testes to any particular tissue in either lineage, a major difference we observe is that functions of *Drosophila* genes broaden over time, whereas functions of mammalian genes narrow over time. Hence, *Drosophila* duplicates eventually acquire broad housekeeping functions, whereas mammalian duplicates acquire tissue-specific functions that may facilitate the evolution of phenotypic diversity across species.

## Methods

### Identification of duplicate and single-copy genes

We downloaded protein sequences, annotation files, and lists of duplicate genes for all genomes from the Ensembl database at http://www.ensembl.org. To obtain a comprehensive list of duplicates in each mammalian genome, we supplemented Ensembl lists with those from the Duplicated Genes Database (DGD) at http://www.dgd.genouest.org and with protein BLAST searches (Altschul et al. 1990), which we performed as previously described (Assis and Bachtrog 2013). Any annotated genes not on these lists were considered to be single-copy genes, and gene families with more than two copies were excluded from our analysis. We quantile-normalized RNA-seq data from mammalian tissues (Brawand et al. 2011) and restricted our analysis to pairs for which both copies are expressed in at least one tissue. Using these expression data, we classified pairs of duplicates as conserved, neofunctionalized, subfunctionalized, or specialized as previously described (Assis and Bachtrog 2013).

### Phylogenetic dating and identification of ancestral single-copy orthologs

We downloaded whole-genome alignments from Ensembl (http://www.ensembl.org) and UCSC Genome Bioinformatics (http://www.genome.ucsc.edu) databases and extracted syntenic regions in all genomes for each duplicate gene. We used parsimony to phylogenetically date the origin of each pair of duplicates. Duplicates that were present in all species or that could not be resolved via parsimony were removed from our analysis. For each pair, the gene copy aligning to the ancestral single-copy gene(s) was considered the parent, and the second copy was considered the child. Orthologs for single-copy genes were also obtained via synteny and aligned with MACSE (Ranwez et al. 2011). PAML (Yang 2007) was used to estimate the *K*_a_ between each pair of single-copy genes.

### Statistical analyses

Mann-Whitney *U* tests were used to compare distributions of relative expression levels, and Fisher’s Exact tests were used to compare observed and expected numbers of genes with highest relative expression in each tissue for conserved and neofunctionalized classes, as well as to compare numbers of genes between classes. All statistical analyses were performed in the R software environment (R Development Core Team 2009).

---

**Figure S1.**
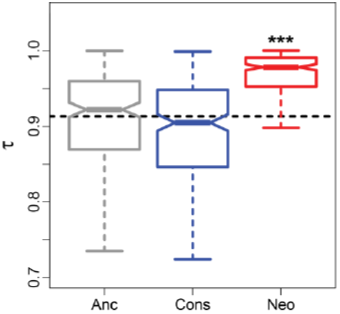
Comparison of tissue-specific expression in conserved and neofunctionalized copies of pairs that underwent neofunctionalization. Boxplots of tissue specificity indices (τ) for ancestral (Anc, gray), conserved (Cons, blue), and neofunctionalized (Neo, red) genes. Dotted black line represents median for single-copy genes, and asterisks show significance relative to distribution of single-copy genes.

